# Combined effect of crop rotation and carabid beetles on weed dynamics in arable fields

**DOI:** 10.1101/2020.12.04.411918

**Authors:** Reto Schmucki, David A. Bohan, Michael J.O. Pocock

## Abstract

Weed management is a resource-intensive practice in arable agriculture, with direct and long-term impacts on both productivity and biodiversity (e.g. plant, pollinators and farmland wildlife). In conventional systems, weed control relies on crop management and herbicide inputs, but for more sustainable production systems, use of herbicides needs to be reduced. This requires a good understanding of the processes that regulate arable weed dynamics in arable fields.

We adopted a systems framework to understand and model interacting components that drive the weed dynamics in 168 arable fields. Within this framework, we built a structural equation model (SEM) to quantify the direct and indirect effects of crop rotation (i.e. crops in the previous three years and the current year) and carabid beetles (Coleoptera: Carabidae) on weed density, seed abundance and accumulation in the seedbank. We included results from a mechanistic approach to infer interactions between seed-feeding carabid beetles and seeds to estimate predation pressure in each field.

Our results show that weeds in arable fields are regulated by crop type, sowing season, and activity density of carabid beetles. We found a direct effect of crop rotation, including both past and current field management practice, on weed abundance in the field and its seedbank. There was also an indirect effect of crops on weed seed accumulation in the seedbank via the effect of seed-eating carabid beetles. The efficiency of weed control by carabid beetles depended on the cumulative predation pressure, which indicates the importance of functional diversity as well as abundance.

Farmers and agronomists can capitalise on the ecosystem services provided by carabid beetles by adapting agronomic practices and crop rotation to maintain a rich fauna of seed-eating carabids in fields and potentially across the agricultural landscapes. When integrated with rotational management practices, this ecosystem services can improve the efficiency of weed management and contribute to the sustainability of cropping systems.

## Introduction

Weed regulation is essential for securing crop yields and quality, as well as for the long-term productivity of arable cropping systems. In conventional agriculture, herbicides and mechanical treatments are the primary tactics used to control competition of weeds with crops. Despite growing concerns about the consequence of the intensive use of agrochemicals on biodiversity (Freemark and Boutin, 1995) and the wider environment (Cross and Edwards-Jones, 2011; Moss, 2008; Silva et al., 2019), the quantity of herbicide use keeps increasing in many regions around the globe (FAO, 2020). Intensive use of herbicides results in important declines of arable plants (Richner et al., 2015; Sutcliffe and Kay, 2000), invertebrates (Bretagnolle and Gaba, 2015; Kraus and Stout, 2019; Pleasants, 2017) and farmland wildlife (Barré et al., 2018; Gibbons et al., 2006; Smart et al., 2000). Such losses in biodiversity undermine the provision of key ecosystem services upon which the productivity and the economy of agroecosystems depend (Bommarco et al., 2013), stressing the need to develop more sustainable approaches that better integrate the benefits of ecological weed regulatory processes.

In many cropping systems, farmers combine several agronomic strategies such as crop diversification, tillage with selective post-emergence herbicides to control weed population outbreaks. This approach offers a cost-effective strategy, but remains highly dependent on synthetic herbicides and is therefore vulnerable to evolution of herbicide resistance (Hicks et al., 2018). Better integration of ecological understanding is urgent, not only to reduce the impact on the environment but also to increase its efficiency and secure long-term productivity of arable systems (Petit et al., 2018). To harness their full potential, integrated weed management approaches must adopt strategies that target different stages of the life cycle to contain their density above ground and their accumulation in the seedbank (MacLaren et al., 2020). By carefully designing crop rotations, farmers can influence weed demography by altering the degree of crop-weed competition, the timing of crop emergence, as well as the density and the height of the vegetation cover. Crop rotations offer many levers to apply diverse selective pressure on weed populations to manage their abundance and composition above ground and in the seedbank (Bohan et al., 2011b; Murphy et al., 2006). Weed seed predation by ground beetles (Coletptera:Carabidea) can reduce seed accumulation in the seedbank and contribute to weeds regulation and cropping systems (Bohan et al., 2011a; Carbonne et al., 2020; Frei et al., 2019; Kulkarni et al., 2015; Petit et al., 2018). Post-dispersal seed predation by carabid beetles is well documented, with studies having focussed on their diet and foraging behaviour (Deroulers and Bretagnolle, 2019; Gaba et al., 2019; Honek et al., 2013; Petit et al., 2014), their distribution in arable fields (Lami et al., 2020), and their response to the surrounding landscape (Petit et al., 2017; Trichard et al., 2014).

Although a growing body of work that shows how crop rotation, agronomic practices and seed predation by carabid beetles contribute to regulating of weeds in arable fields, their effects must be considered as part of a comprehensive system (Hawes et al., 2016) where weed dynamics are influenced by both agronomic and ecological factors. Such approaches are also necessary to identify tradeoffs between the costs of arable weeds, and their benefits, e.g. in supporting regulating services (Bretagnolle and Gaba, 2015), and to adequately measure the direct and indirect impacts of specific cropping systems and integrated management practices on weed regulation (Mézière et al., 2015).

Here we investigate the direct and indirect effects of crop rotation, field management and seed predation on the density and the dynamics of arable weed communities. We built a conceptual model that explicitly integrates the action of both agronomic and ecological drivers on weed dynamics and at each stage of their life cycle (*see Structural Equation Model* in *Methods*) and is based on theoretical knowledge and results obtained from previous investigations. Using structural equation models (SEM), we then tested this conceptual model against empirical data. One benefit of using SEMs is their capacity to explicitly to account for the temporal sequence that underpins the different events. In our model, we aligned the chronology of the events to investigate how factors connect and affect the abundance of weed throughout the life cycle in arable cropping systems. Using an existing data set from a large-scale field sampling of carabids and weed in arable cropping systems, we analyse the direct and indirect effects of 1) crop rotation; 2) agronomic practices; and, 3) the density of carabid beetles on the density of arable weeds growing and stored in the seedbank of arable fields. With this model, we can disentangle part of the complexity of integrated weed management by measuring the specific and combined effects of crop rotation, agronomic practices and seed predation by carabid beetles on weed abundance in arable fields. This quantitative understanding is useful for developing sustainable weed management strategies that do not compromise crop yield, ensure long-term productivity and minimise the impact on biodiversity and the environment.

## Methods

In our study, we used an extant dataset of ecological surveys in arable fields collected through the Farm Scale Evaluations (FSEs) of genetically modified, herbicide-tolerant (GMHT) crops (Firbank et al., 2003). The FSE field trials were established in 2000 and ran until 2002 to assess the potential impact of adopting genetically-modified crop on arable biodiversity within and surrounding agricultural fields (Hails, 2000; Watkinson et al., 2000). The 256 fields included in the FSE study were carefully selected to provide a representative sample of the British arable farming system (Champion et al., 2003; Firbank et al., 2003). The trials focus on four crops, namely spring-sown maize, spring- and winter-sown oilseed rape and a spring-sown beet (forage and sugar beet). In the trials, each field was split into two halves with the GMHT variety sown on one side and the conventional variety on the other to contrast the responses observed in genetically modified and conventional crop systems. Here, however, we restricted our analysis to the data collected in the conventional part of the fields. During the FSE field trials, farmers were audited to document the crop management practices, including herbicide treatments used in each field which was based on farmers’ decisions and what they considered consistent with cost-effectiveness (Champion et al., 2003; Firbank et al., 2003). In order to document how herbicide usage and concentration varied across crops, we described the list of active ingredients, their concentration and timing of their application (Figure S1).

In each field, data were collected along 12 parallel transects arrange perpendicularly along three edges of the fields and ran into the crop. Sampling points were set along each transect and located at 2, 4, 8, 16 and 32 m from the field margin (see full detail in Firbank et al., 2003). Because we were interested in weed management in the crop field, we excluded data collected near the edge (2 and 4 m) and focused on sampling points located at 8, 16 and 32 m. Data collected in the field trials can be retrieved from NERC Environmental Information Data Centre (Scott et al., 2012a, 2012b, 2012c, 2012d).

### (a) Seedbank

Seed density contained in the seedbank of each field was assessed before the crop was sown (t_0_), and before the crop was sown in the following year (t_1_). In each field, soil samples were systematically collected along four of the 12 transects before the sowing of the crop to establish a baseline of the seedbank at t_0_ (Heard et al., 2003b). Size and composition of the seedbank were obtained from 1.5 kg soil samples, taken to a depth of 15 cm at 2 and 32 m along four transects per field. Soil samples were weighed and passed through a 10 mm mesh sieve before being placed in a plastic tray to a depth of 4 cm and arranged in an unheated glasshouse on capillary matting ke pt moist. Seedlings that emerged from the seed bank samples were identified and removed from the trays for 18 weeks after sample preparation. The following year, soils samples were collected at the same locations, at nearly at the same time of the year, to establish the size and composition of the seedbanks at t_1_ (Heard et al., 2003b).

### (b) Standing weeds

The composition and density of the above-ground weed community were sampled at two times in the season along each of the 12 transects (see Perry et al., 2003). Individual standing weed counts were made once after the sowing of crops and once before harvest after all treatments in 0.25 × 0.5 m quadrats set each sampling points along the transect (Heard et al., 2003b). These counts were used to estimate the density (individuals m^-2^) of standing weed observed in each field beyond 8 m from the edge.

### (c) Seed rain

In each field, a measure of the annually-produced and dispersed seeds was obtained by sampling the seed rain, using four seed traps set at ground level and monitored repeatedly over the growing season (Heard et al., 2003b). Traps of 10 cm in diameter and 10cm high where sunk into the ground at 2 and 32 m on four transects. Seeds were collected fortnightly, starting after first anthesis of weeds and until crop harvest. The seeds collected over the crop season were pooled, identified to species and classified as ‘viable’ and ‘non-viable’ based on their physical attributes (see Heard et al. 2003; Bohan et al. 2005). For our analysis, we quantify the total seed rain as the sum of viable seeds sampled at 32 m over a year.

### (d) Carabid beetle activity-density

Soil-surface active invertebrates were sampled from pitfall traps (6 cm diameter) placed at 2, 8 and 32 m along four of the 12 transects. Pitfall traps were open for two-week, on three sampling occasions during the year (Bohan et al., 2005; Brooks et al., 2003). All carabids collected in the pitfall trap were counted and identified to species, and then summed across the year to create a year total for analysis. For consistency in the sampling effort and season between spring and winter-sown crops, we only considered carabid counts collected at in July and August for spring crops and April - May and June-July for the winter crop, excluding samples collected in September and October in winter-sown oilseed rape (Firbank et al., 2003). For each field, we estimated the year-total activity-density of granivorous (e.g., *Harpalus* and *Amara* spp.) and omnivorous (e.g., *Pterostichus* and *Bembidion* spp.) species from the pooled counts sampled in the eight pitfalls (i.e., 8 and 32 m set along four transects). Because we were interested in estimating the effect of seed predation on weed management, we focused on carabid beetle species that were capable of consuming weed seeds and excluded species that are known to be obligatory carnivores (Luff, 2017).

### (e) Carabid predation pressure

We quantified predation pressure as the number of seeds consumed by the assemblage of carabids present, taking into account carabid seed preferences, the community of seeds present, and carabid energetic requirements (Pocock et al., 2020). In their work, Pocock et al., (2020) derive a predation pressure index from an inferred carabid-seed food web. In this food web, the interactions were inferred based on species frequency-dependent foraging and size-dependent preferences obtained from the literature (Gendron, 1987; Honek et al., 2013; Petit et al., 2014), and the interaction strengths were calculated by scaling up by carabid density and energetic requirements.

### (f) Crop sequence and rotation history

After the FES trials, a follow-up study was conducted by Bohan et al. (2011b) to document and investigate the effect of crop rotation on weed management. Participating farmers were contacted and asked about the crop history that was grown in for each field, up to nine years before the FSE trials. Farmers provided this information, and the detailed crop sequence was available in 168 fields. From this sequence, we derived two indices to characterise the type of crop and their continuousness (pattern along the sequence) in the cropping systems. We calculated a “cereal index” to capture the frequency and pattern of cereal crops grown in the three years prior to the FSE trials. We also calculated a “winter index” to quantify the frequency and pattern of winter sowi ng crops grown within the three years. Each index range from 0 to 7 and quantify the pattern in the crop sequence, based on both consecutiveness and recentness of the focal state, where 0 indicates the absence of the focal state (cereal or winter-sown crop) and 7 indicates a continuous sequence over the three years (see Figure S2).

### Structural Equation Model

In order to obtain a system-based understanding of the interacting components, including the effect of both agronomic and ecological factors, we built our conceptual model around the life cycle of arable weeds growing in conventional cropping systems (Figure 1). In our model, we used theoretical knowledge and results of previous investigations to integrate relationships that hypothesises how the different factors affected each development stages and the dynamics of arable weeds in arable fields. Our model accounts for the temporal sequence of the weed life cycle, as well as the sequence (precedence) of events that underpin specific cropping systems (crop rotation history, field management) and field environment (carabid abundance). In our model, we aligned the chronology to investigate how factors connect and affect the initial seedbank (t_0_), the abundance of standing weeds, the seed rain and the activity-density of carabid beetle in the field, their predation pressure and their combined effect on the seedbank in the following year (t_1_).

**Figure 1.**
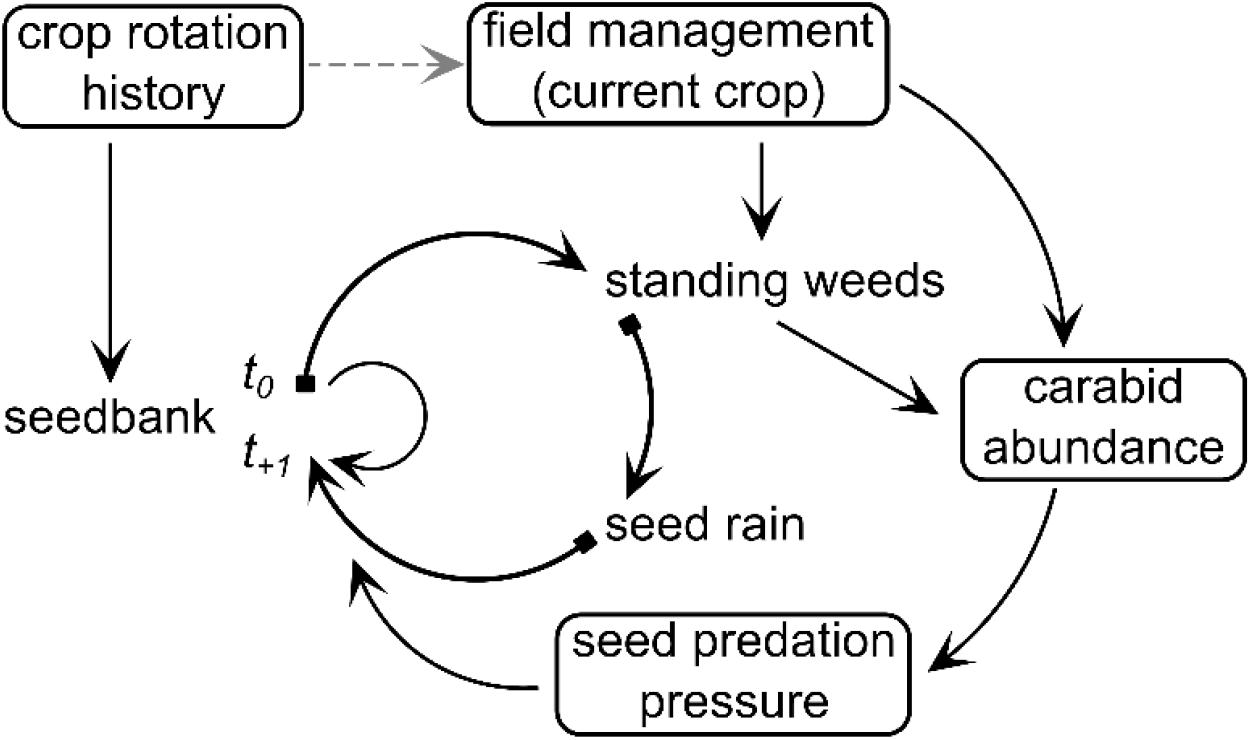
Graphical conceptual model that represent the drivers and their interactions with the life cycle of arable weeds in crop fields. This model integrates both agronomic (crop history and field management) and ecological (seed predation by carabid beetles) components, as well as the chronology (direction) of their direct and indirect effect on the dynamics of arable weeds stored in the seedbank and growing in crop fields.

In our model (Figure 1), we integrated the effect of crop rotation on the seedbank at t_0_ as an effect of past field and crop management (tillage, herbicide option and sowing season). We specifically focused on the crop type (cereal versus non-cereal), and the sowing-season (spring and winter) as these two aspects affect management and herbicide regimes, as well as the phenology of the crop cover (Bohan et al., 2011b; Heard et al., 2003a).

Based on our conceptual model (Figure 1), we constructed a Structural Equation Model (SEM). In our SEM, the recent crop history documented in each field was summarised using their three-year cereal and winter sowing indices (see Section *f* in Methods above). To account for the potential non-linear effect of crop sequence on the seedbank, we added a quadratic effect between each index and the seedbank t_0_. The SEM also included the effect of crop management with the variable *field management (current crop)* which was coded with three binary dummy variables for beet, maize and winter oilseed rape and setting spring oilseed rape as the reference value. With the current crop, we aimed to capture the effect of specific crop and their management on the density of standing weed. We expect the specific sowing season and herbicide usage (Figure S1) associated to the different crops to affect the germination, the establishment and the growth of weeds and thereby influence the density of standing weed, together with the size of the seed pool contained in the seedbank at t_0_. Crop management is not only expected to influence weed but also the activity-density and the composition of carabid beetle foraging in arable fields through its effect on the stability of resting and foraging habitat (Menalled et al., 2007; Trichard et al., 2014). Activity-density of carabid beetle is also expected to be influenced by the density of standing weed density as they partly determine the quality and quantity of resources available to phytophagous species foraging in arable fields (Frei et al., 2019). Finally, we added four effects to predict the abundance of seeds in the seedbank in the following year (t_1_). Firstly, the effect of the starting seedbank abundance at t_0_, which accounts for seeds that remain dormant over the cropping season (Mahé et al., 2020). Secondly, the seed rain, which accounts for seeds measured to be shed by the standing weeds. Thirdly, a direct effect linking the standing weed with the seedbank t_1_, which accounts for seeds shed by standing weeds, but not captured in the seed rain sample. This structural part (direct effect) was added after examination of the remaining covariance when the model only included the effect of the seed rain, i.e. the indirect link between standing weed and seedbank t_1_. Fourthly, the effect of carabid beetles on the seedbank at t_1_ by including the predation pressure index derived from the inferred carabid-seed food web in each field (see details in Pocock et al., 2020). With this last relationship, we accounts for the effectiveness of carabids in regulating arable weeds, based on their dietary preference and calorific requirement which are expected to have important implication in carabid seed consumption and contribution to weed management (Gaba et al., 2019; Honek et al., 2013; Petit et al., 2014).

We specified the SEM and the relationships between the endogenous and exogenous variable as follow:

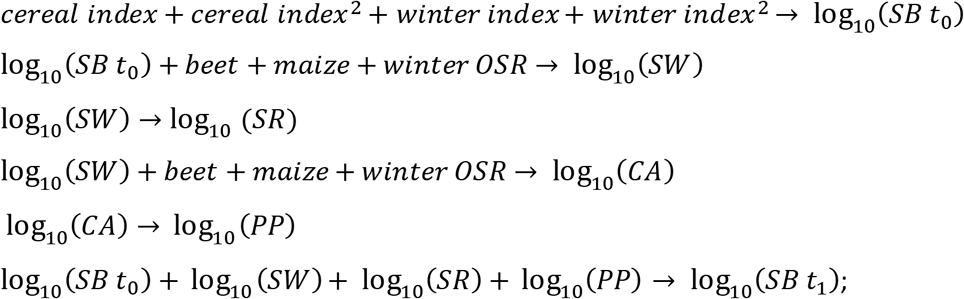

where *SB* is weed count in the seedbank, *SW* in the standing weed, *SR* in the seed rain, *CA* is the active-density of carabid beetles, *PP* the predation pressure and *OSR* stand for oilseed rape.

### Statistical modelling

Fitting complex SEMs can be challenging and it requires large data sets to reach convergence and produce reliable confidence intervals. However, recent computational advances have led to the development of Bayesian approaches for SEMs (Merkle et al., 2020) that allow for the fitting of complex models because they address issues of non-convergence and improve estimations of parameter confidence intervals (Smid et al., 2020). We, therefore, fitted our SEM within a Bayesian framework, using the MCMC sampler Stan 2.21-0 (Carpenter et al., 2017), in combination with the R packages lavaan 0.6-7 (Rosseel, 2012) and blavaan 0.3-10 (Merkle and Rosseel, 2018). The estimates for the Bayesian SEM were derived from four chains with 10,000 samples, after 5,000 burn-in iterations and a thinning factor of one. When fitting our model, we used weakly informative prior for the beta parameters (regression coefficient), using a normal distribution with mean 0 and standard deviation of five. This prior was preferred to the default normal(0, 10) available in blavaan (Merkle and Rosseel, 2018) because we used the log_10_ transformation for each variable derived from counts (seedbank, standing weed, seed rain, carabid activity-density, and predation pressure). Therefore, coefficients are expressed on a log-log scale and must be interpreted as expected proportional differences per proportional change (i.e. per cent of the difference of *y* for a per cent change in *x*). Note however that rotation indices and crops were not log-transformed, and the coefficients to the transformed count must be interpreted as multiplicative differences (i.e., *y* = 10^coefficient^ for a change of 1 in *x*). Because our priors contain little information, our posterior estimates are dominated by the information available in the data. We assessed the MCMC chains convergence through visual observation and the potential scale reduction factor (*Rhat* ≈ 1). We estimated the validity of our conceptual model from the overall fit judged from the posterior predictive *p-*value (*ppp*). For each endogenous variables (dependent), we calculated the proportion of variance explained from the Bayesian *R*^2^ derived from the standardised variance estimates. Based on the results of our SEM, we implemented a multivariate response model(Path model) within a Bayesian framework, using the package brms 2.13-5 (Bürkner, 2018) to define our model with the same prior as specified in the SEM, normal(0, 5), with four MCMC chains, 10,000 samples and 5,000 burn-in iterations and a thinning factor of one. From the posterior samples, we estimated the conditional effects for each local model included in the SEM(Figure 1).

## Results

The system framework provided by the structural equation model (SEM) was useful to integrate and identify the effect sizes of the different drivers shaping the structuring the abundance and dynamics of arable weeds in cropping systems (Figure 2). Our SEM, build around the life cycle of arable weeds, enabled us to integrate both agronomic and ecological factors and explained 43 percent of the variation observed in the density of weed contained in the seedbank after one crop season (Table 1). While past field cropping history explained five percent of the variance of contained in the seedbank at *t*_*0*_, when we added the components of the weed life cycle, crop management and the regulatory effect provided by carabid beetles, the predictive power of our model was substantially increased, with R^2^ for local models ranging from 0.05 to 0.63 (Table 1).

**Table 1.**
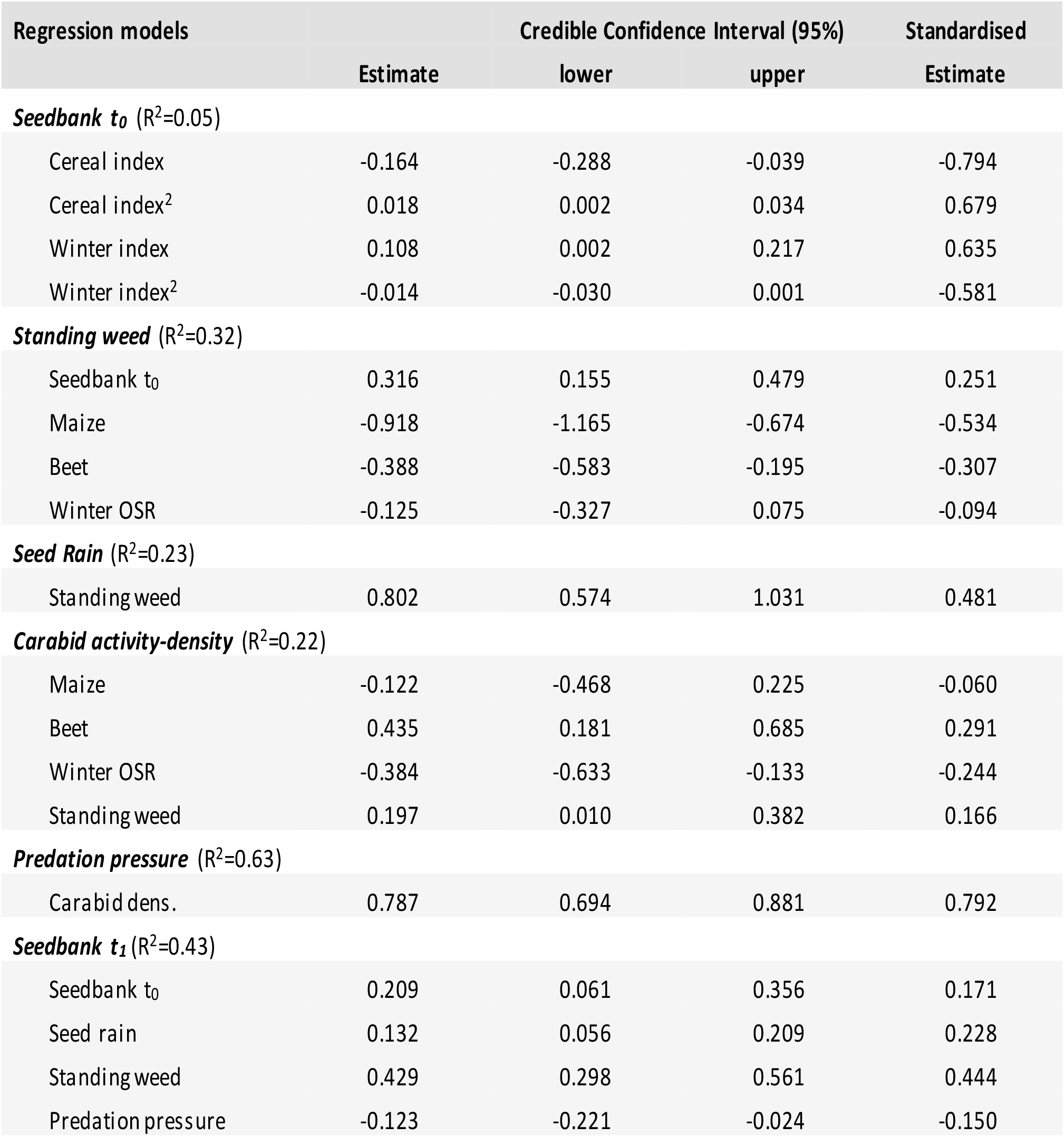
Detailed SEM estimates and 95% credible confidence intervals and R^2^ for each endogenous variables. The SEM was fitted with Stan MCMC sampler, using 10,000 samples and 5,000 burn-in iterations. The global model offers an acceptable fit to the data (Bayesian fit: n = 168, ppp = 0.086).

**Figure 2.**
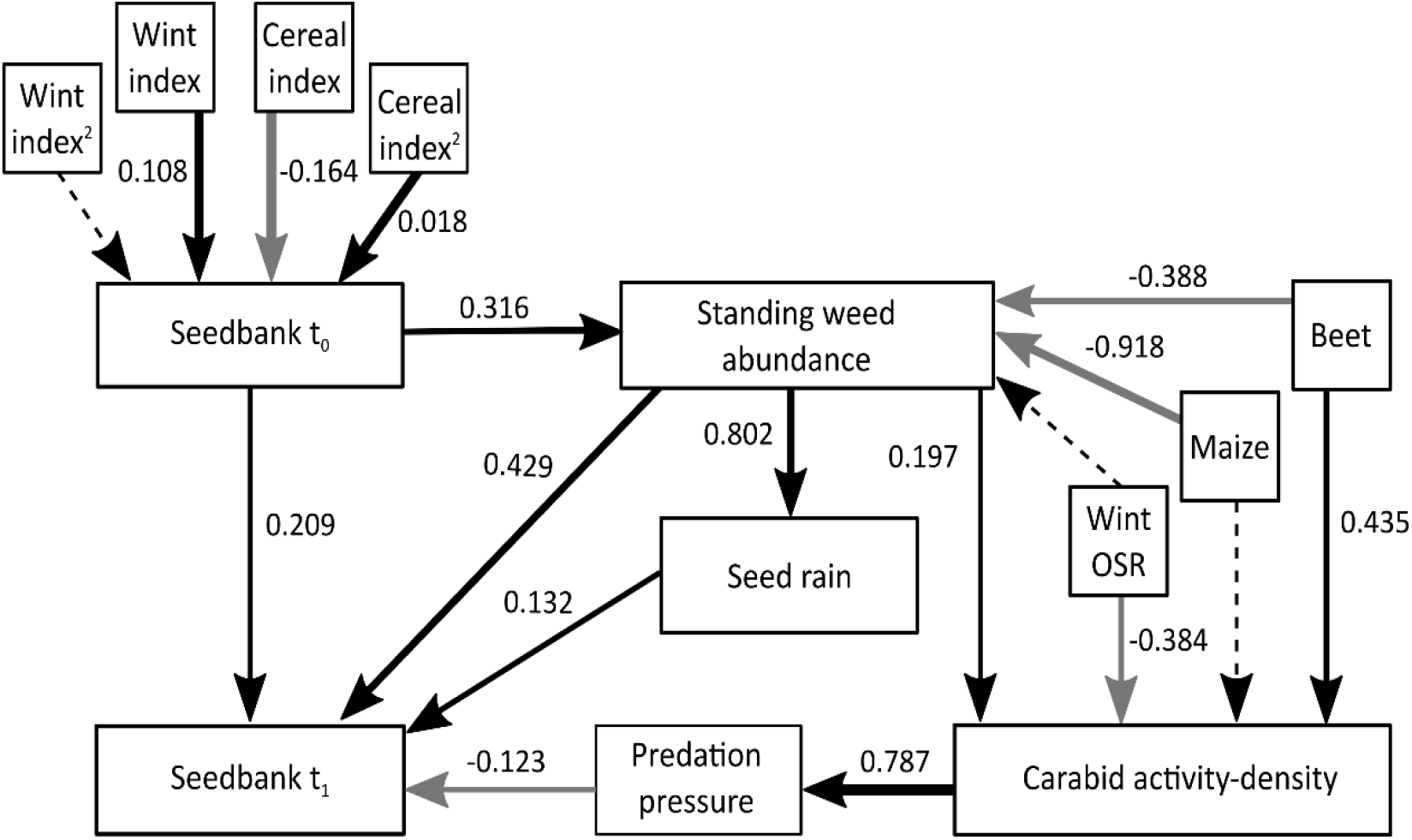
Summary of the structural equation model (SEM), with unstandardised estimates of the effects of crop rotation, field management and predation pressure of seed-eating carabid beetles on the density of weeds growing in the fields and stored in the seedbanks at t_0_ and t_1_. Solid arrows represent significant effects, black for positive and grey for negative, and dashed arrows display non-significant effects. The thickness of each arrow is relative to the standardised effect size. The global model provides an acceptable fit to the data (Bayesian fit: n = 168, PPP = 0.086).

In our study, we focus on the impact of the structure of past crop rotations (crop type and seasonality), the effect of crop management crop and the influence of foraging ground beetles feeding on weed seeds. Our SEM enabled us to account for the temporal aspect of each relationship, identifying both the direct and indirect effects of the drivers and showing how regulation services provided by carabid beetle are intertwined within the dynamics of the weed and cropping system (Figure 2). Our model shows the contribution of the different components of the system that regulate the abundance of seeds growing in the field and accumulating in the seedbank over time. Interestingly, our results suggest that predation pressure provided by carabid beetles contribute significantly to reduce the accumulation and the size of the seedbank, having nearly the same effect size, but in the opposite direction as the seed rain (Table 1).

Although the crop history (rotation) defined by the cereal and winter-sowing indices had a relatively low explanatory power, the model shows that the abundance of seed contained in the seedbank at *t*_*0*_ was significantly lower in fields where recent crop sequence had non-consecutive cereal (including a break crop) (Figure 3a). The quadratic effect of the cereal component of the rotation shows its importance for regulating the size of seedbank over time, as well as the density the weed growing in the field and competing with crop. Interestingly, fields that had recent winter sowing crop (winter index level 3 and 4) tended to have slightly larger seedbank. Nevertheless, this effect was relatively small and it is unclear if this effect persists as winter crops are sown over multiple years (Figure 3b) (Table 1).

**Figure 3.**
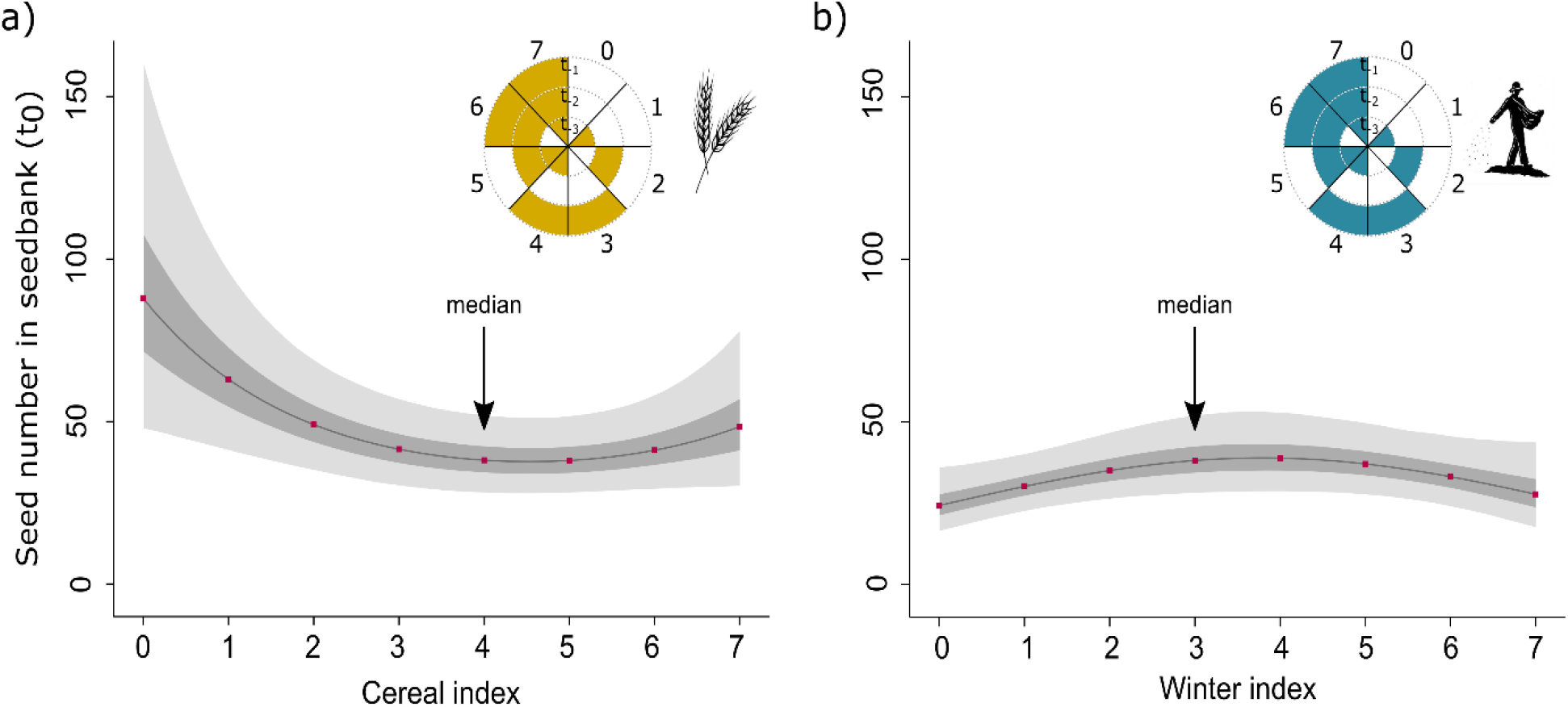
Conditional effects for a) cereal and b) winter sowing crop on the number of seeds in the seed bank at t_0_. Conditional effects were derived from a Bayesian multivariate response model with seed density log_10_ transformed and where all other predictor variables set to their mean. Indices are derived from the 3-year crop history (three years before t_0_) and range from 0 to 7, where 0 indicate the absence of cereal or winter sowing and 7 three consecutive years of cereal or winter sowing (Figure S2). The 50% and 95% credible intervals are depicted by the dark and light grey areas around the point estimates. For each index, we identified the median value of the index with an arrow.

Identity of the crop grown in arable fields and the associated management practices, including herbicide regimes (Figure S1), and potentially crop-weed competition had a substantial effect on the density of weed growing in the field, with maize and beet having markedly fewer weeds than oilseed rape (Figure 4a). Fields with winter-sown oilseed rape tended to have a lower density of weeds than spring-sown oilseed rape (Figure 4a). This difference suggests that sowing season and the phenology of the crop (canopy development) impact weed growth through its effect on weed-crop competition. Although the density of standing weeds growing in crop field had a positive effect on the activity-density of carabid beetles, it was clear that other factors influenced the density of carabids because their abundance was substantially higher in fields growing beet, even though this crop did not have the highest density of arable weeds (Figure 4).

**Figure 4.**
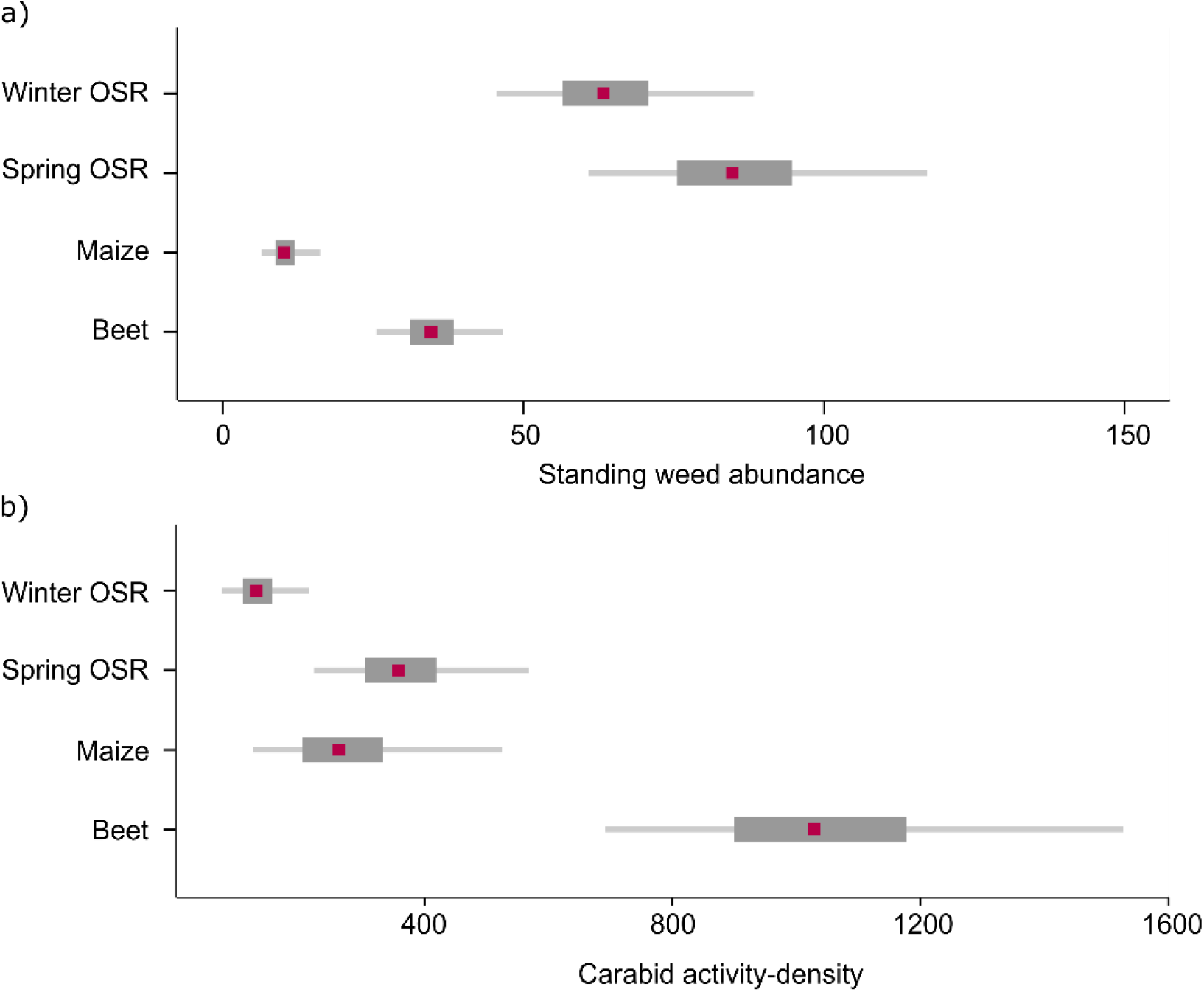
Conditional effect of crop sown at t_0_ on a) standing weed abundance and b) carabid activity-density. *Conditional effects were calculated with all other predictor variables set to their mean*. The 50% and 95% credible intervals are depicted by the dark and light grey areas around the point estimates. Estimates are derived from a Bayesian multivariate response model, with the count of standing weed abundance and carabid-activity density log_10_-transformed. OSR = oilseed rape.

Overall, our models show that the density of standing weeds growing in the field had the largest effect on the size of the seedbank in the following year (Figure 5b). However, our results also show that the size of the seedbank is determined, at least partly, by the activity of seed-eating carabids beetles (Figure 5c). This suggests that when crop rotation are carefully designed, and in-field conditions are managed to increase the density and the predation pressure exerted by carabid beetles, their combined effects can help regulate the size of the seedbank and over time help managing the density of weeds growing in the arable fields (Figure 5). The increase of the activity density of seed-eating carabid beetles observed when the weeds density in crop field increases suggest that the regulation service provided by these invertebrates is likely to vary with the need for weed management (Figure 2).

**Figure 5.**
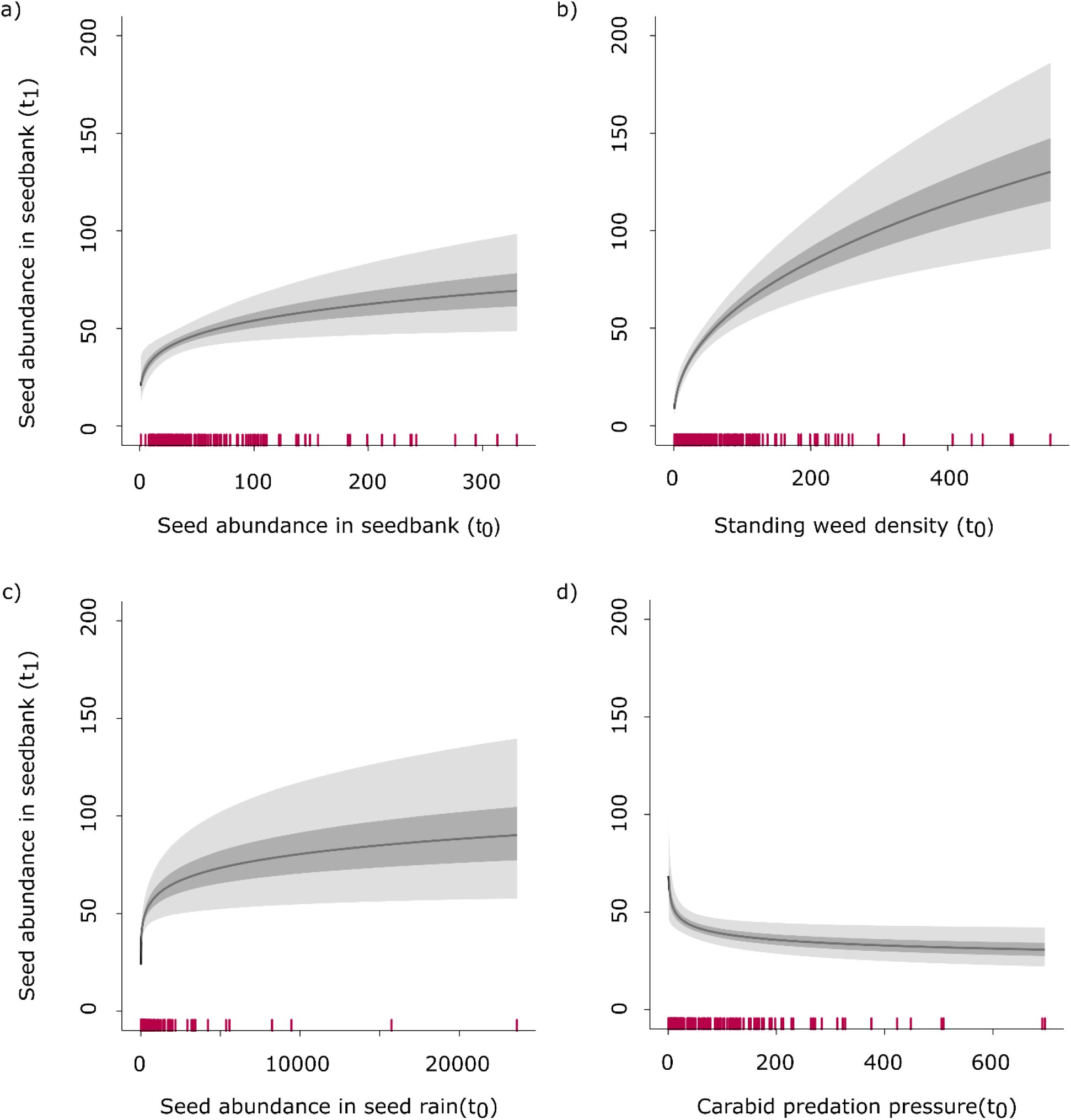
Conditional effects of a) the effect of seedbank density at t_0_, b) standing weed density per square meter at t_0_, c) the number of seed in seed rain on seedbank density at t_+1_ and d) the predation pressure of seed-eating carabid on seed density in the seedbank at t_1_. Conditional effects were calculated with all other predictor variables set to their mean. The 50% and 95% credible intervals are depicted by the dark and light grey areas around the mean estimates. Estimates are derived from a Bayesian multivariate response model, with count variables being log_10_ transformed.

## Discussion

Sustainable weed management is essential for maintaining the productivity of cropping system while minimising the negative impact on the environment. Development of strategies that integrate both agronomic and ecological processes into cropping systems will contribute to ensure food security and support essential biodiversity that agroecosystems depend on. In this study, we clearly show that agronomic practices associated to cropping systems and the regulation services provided by carabid beetles are intertwined and together influence the density and the dynamics of weeds stored and growing in arable fields. Such understanding can provide new perspectives that will help farmers and agronomists integrate nature-based solutions to maintain high yield while reducing the economic and environmental cost of weed management.

Integrated weed management applied in conventional agricultural systems has always aimed at combining multiple approaches to increase the efficiency of weed management. However, the complexity and the level of integration of ecological processes regulating the density and controlling for weed outbreaks is often very limited (MacLaren et al., 2020; Petit et al., 2018). Because seeds in the seedbank are buried in the soil and remain invisible to farmers, it is challenging to explicitly accounted for in weed management plans. This lack of information has probably contributed to the fact that weed management is mainly applied responsively where the density of arable weeds is controlled through herbicides application. This approach is effective and influences the density of weed accumulating in the seedbank, but because it does not explicitly account for the regulation services provided by biological agents such as seed-eating carabids, farmers cannot fully integrate the cost and benefits of such approaches. By systematically accounting for the effect of crop rotation and seed predation on the seedbank, and how these effects cascade across the different life stages of arable weeds, we can reveal areas in the weed management plans where synergy can be created and where tradeoffs must be carefully considered (Mézière et al., 2015). Such information is difficult to derive from independent analysis accounting for the action of each aspect separately.

We show that farmers can actively reduce the abundance of weed contained in the seedbank by managing the frequency and consecutiveness of cereal crops and, but to a lesser extent, by varying the sowing season. This result confirms the well-known benefits of including ‘break crops’ in rotation, changing the selective pressure applied on the weed community and thereby regulating the demography of different species or type of arable weeds (monocot versus dicot) (Derksen et al., 2002). By changing the sowing season, producers are not only affecting the timing of soil disturbance regimes (tillage and drilling), but also the phenology of the crop cover and thereby changing crop-weed competition (Colbach et al., 2014). The competitive of taller crops suppress the germination and growth of weeds, but also reduce reproduction success and the number of seeds re-entering the seedbank.

Weed regulation through the effect of carabid beetles has been documented in many studies, and our results confirm how these biological agents can contribute and strengthen the efficiency of integrated weed management approaches (Bohan et al., 2011a; Kulkarni et al., 2015). Because field management and the density of standing weeds in arable fields influence the number of seed-eating carabid beetles foraging in the field, optimal weed management would aim at regulating rather than eradicating weeds. This would allow supporting effective level of predation pressure in the field in order to limit the number of seed re-entering the seedbank. Weed regulation through the action of natural predators not only reduces the number of weeds in the seedbank but can also reduce the use of synthetic herbicides and the risk of evolution of herbicide-resistance. Although carabid beetles do have preferences in their seed consumption (Honek et al., 2007; Petit et al., 2014), carabid diet is relatively broad and largely driven by seed:carabid size (Pocock et al. 2020). Therefore, the regulation service provided by carabid beetles is likely to increase taxonomic and functional diversity and control the dominance of highly competitive weeds.

When considered together, appropriate crop rotations and effective seed-predation provided by seed-eating beetles provide tools additional to synthetic herbicides to manage weeds in arable fields and their effect on crop yield. By using a system framework that accounts for the effect of different drivers regulating the size and the composition of weed communities, we give farmers the means to improve the sustainability and resilience of agroecosystems. Weed management strategies that integrate and connect agronomic and ecological regulation components can reduce the reliance on synthetic herbicides and promote the long-term provision of highly valuable ecosystem services to secure the future of food production systems.

## Data accessibility

The FSE data, including the farm management data, are available from the NERC Environmental Information Data Centre (https://catalogue.ceh.ac.uk/documents/876358e4-62f7-4386-99e1-7d3eac223e03).

## CRediT author statement

**RS:** Conceptualisation, Methodology, Data Curation, Formal Analysis Writing – Original Draft, Visualisation, Writing – Review and Editing; **DAB:** Funding acquisition, Project administration, Conceptualisation, Methodology, Writing – Review and Editing; **MJOP:** Funding acquisition, Project administration, Conceptualisation, Methodology, Writing – Review and Editing

## Acknowledgments

**RS** and **MJOP** were funded by Defra (contract SCF0313). **DAB** was funded by the Agence Nationale de la Recherche (ANR). This work is an output of the PREAR project (Predicting and enhancing the Resilience of European Agro-ecosystems to environmental change using crop Rotations), a partnership between INRA, UKCEH, University of Copenhagen and Aarhus, Solagro and Szent István University. The PREAR project was funded as part of the European FACCE SURPLUS (sustainable and resilient agriculture for food and non-food systems) and the ERA-NET co-fund scheme under Horizon 2020 programme formed in collaboration between the European Commission and a partnership of 15 countries in the frame of the Joint Programming Initiative on Agriculture, Food Security and Climate Change (FACCE-JPI).

## Supplementary material

**Figure S1.**
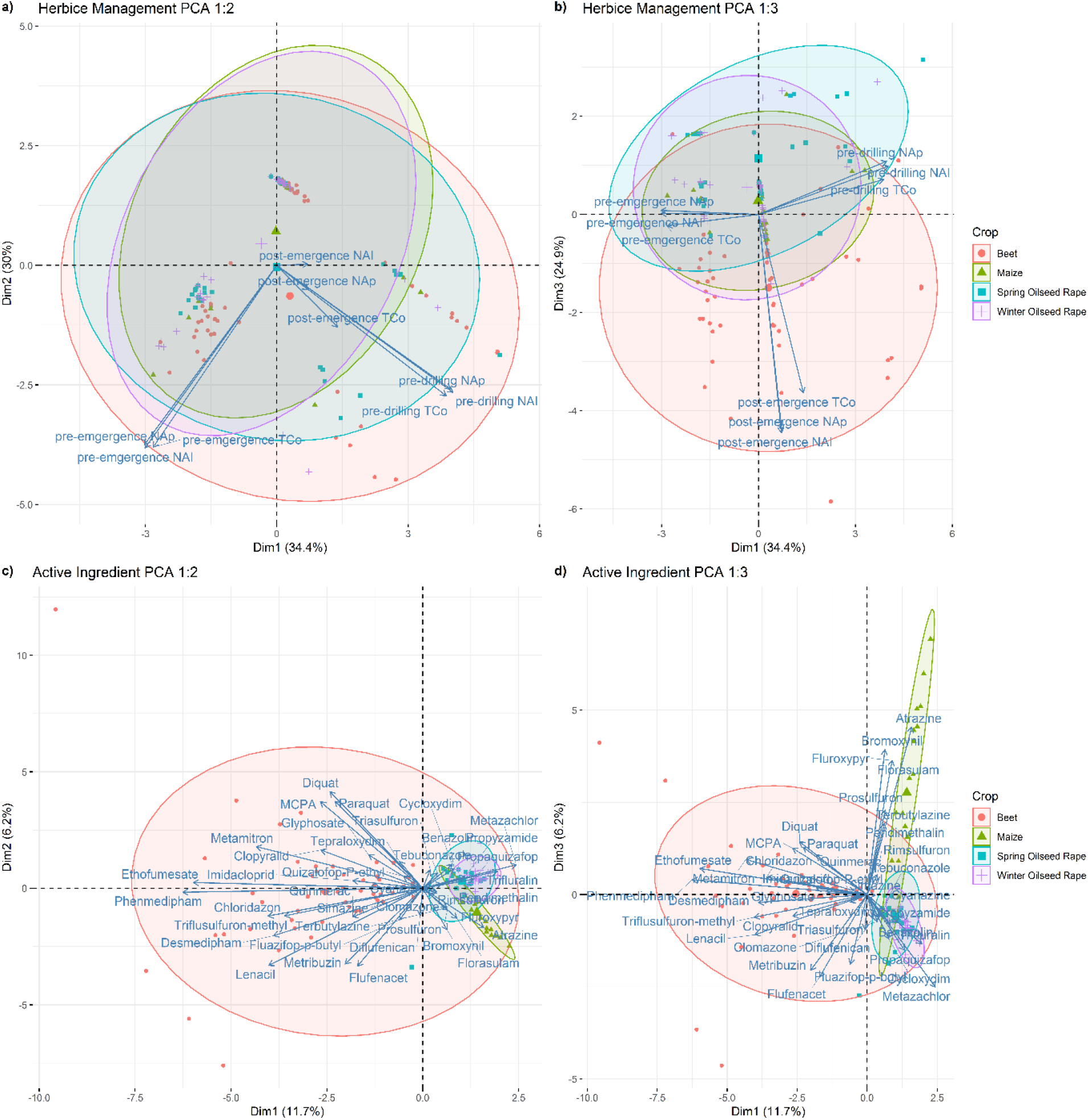
Principal Component Analysis (PCA) results with a) the first and second components and b) the first and third components of the total amount of herbicide applied per hectare (TCo), the number of application (Nap) and the number of active ingredients (NAI) applied on four conventional crops (spring-sown beet, spring-sown maize, spring-sown oilseed rape and winter-sown oilseed rape) grown across 168 fields monitored during the Farm Scale Evaluation field trials. c)The first and second components and d) the first and third components of the PCA computed on active ingredients (frequency and diversity) used in four crops across 168 fields.

**Figure S2.**
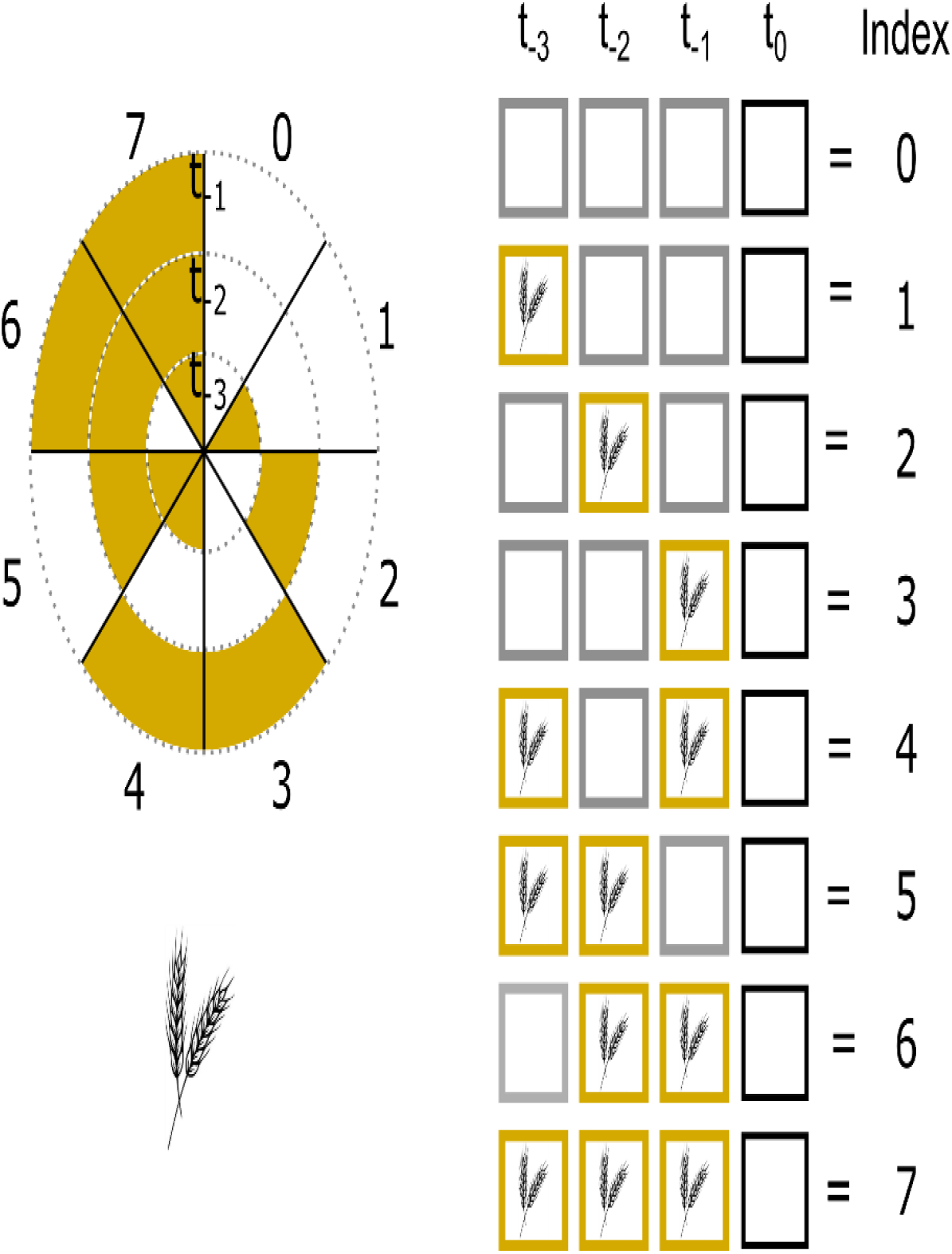
Schematic representation of the index used to integrate the cereal (illustrate above) and the winter-sown component contained in the crop sequence (rotation) observed in the three years preceding the sampling of the reference seedbank (*t*_*0*_) in the Farm Scale Evaluation field trials. Ranging from 0 to 7, the index represents the change in consecutiveness and recentness of the focal state (cereal crop or winter-sown crop), where 0 indicates the absence of the focal state and 7 indicates a continuous sequence over the 3-year period.

